# Evolutionary Origins and Synthetic Reconstruction of Gingerols

**DOI:** 10.64898/2026.07.28.741391

**Authors:** Weihao Wang, Qingmin Li, Jun Zhang, Xiaosong Yu, Xinyi Li, Fanze Meng, Haiyang Jia, Bing Hu, Bo Lv, Lei Qin, Chun Li

## Abstract

How new enzymatic activities become integrated with existing metabolic networks to generate lineage-restricted plant chemistry remains a central question in metabolic evolution, exemplified by gingerol biosynthesis in *Zingiberaceae*, whose complete enzymatic basis has remained elusive. Here, we resolve this problem by elucidating the evolutionary origins and the full biosynthetic pathway of gingerol through single-cell transcriptomics, functional screening, structural biology, comparative genomics and pathway reconstruction. A coordinated metabolic program operates in ginger oil cells, thereby guiding the identification of diketide-CoA synthases, gingerol synthases (GS) and gingerol reductases that together constitute the complete pathway. GS remodels the conserved type III polyketide synthase scaffold to organize a fatty acyl-CoA and a phenylpropanoid-derived intermediate for gingerol formation. Furthermore, GS arose through duplication and limited active-site remodeling of a curcumin synthase-like enzyme, accompanied by coordinated recruitment of precursor-supply and reduction pathways. Translating these principles into yeast achieves the first de novo microbial production of 6-gingerol from glucose at 12.4 mg/L. These findings reveal the evolutionary logic by which plants recruit and coordinate novel enzymatic activities to establish lineage-restricted metabolic program, providing a framework for understanding and engineering specialized metabolism across the plant kingdom.

## Introduction

The structural diversity of plant specialized metabolites is a product of long-term evolution^1–3^, yet the mechanisms by which closely related species remodel shared ancestral pathways to generate distinct chemical phenotypes remain poorly understood^4–6^. This contrast is particularly striking in the *Zingiberaceae*, where curcumin and gingerol share phenylpropanoid-derived aromatic precursors but are built through fundamentally different carbon-skeleton assembly logics^7^. Gingerols are the principal bioactive constituents of ginger, contributing substantially to its nutritional and medicinal value^8^. Curcumin is formed through sequential condensations that generate a diarylheptanoid scaffold, whereas gingerol biosynthesis requires the coupling of a fatty acyl chain with a phenylpropanoid-derived polyketide intermediate^9^. Although the key enzymes involved in curcumin biosynthesis have been identified^10–11^, the core enzymes responsible for gingerol scaffold assembly and reduction have remained unknown^12–14^. This knowledge gap not only hinders our understanding of the origins of lineage-specific metabolic innovation in the genus *Zingiber*, but also constrains heterologous production of these valuable compounds through synthetic biology approaches.

The contrasting carbon skeletons of curcumin and gingerol likely reflect functional divergence of type III polyketide synthases (PKSs), which use a conserved Cys-His-Asn catalytic triad to assemble diverse scaffolds from different starter and extender units^15^. The well-characterized chalcone synthase (CHS) is broadly involved in flavonoid biosynthesis, whereas diketide-CoA synthase (DCS) and curcumin synthase (CURS) represent a functional divergence of this family within the *Zingiberales*^16–17^. However, formation of the gingerol skeleton requires a previously uncharacterized PKS, which could couple a linear fatty acyl-CoA with feruloyl-β-keto acid through decarboxylative Claisen condensation^7,9^ (Fig 1a and b). This uncharacterized PKS might have evolved from related enzymes through local remodeling of the active site. Identifying this missing enzyme and elucidating its catalytic mechanism therefore constitute a critical step toward understanding the biosynthetic origin of gingerols.

**Fig. 1.**
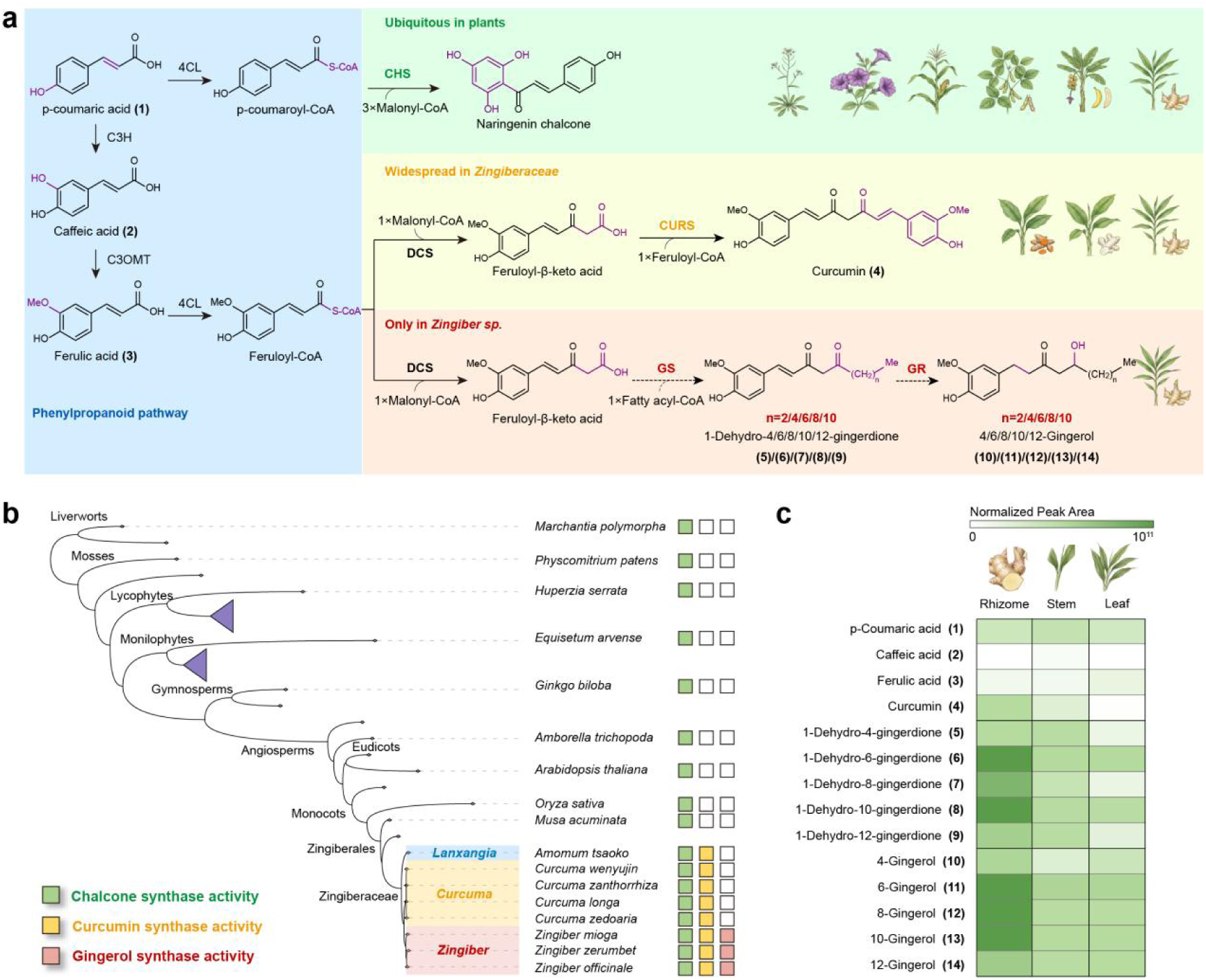
Phylogenetic restriction and organ specialization of gingerol biosynthesis. **a)** Proposed biosynthetic framework linking the broadly conserved phenylpropanoid pathway to three type III polyketide-derived chemical traits with progressively restricted phylogenetic distributions. Dash arrows represent uncharacterized step, black arrows represent characterized steps, and the overlapping arrows represent a multi-step reaction. Compound numbers correspond to those used in **c**. All plant illustrations appearing in the picture were generated by Gemini Banana2. **b)** Phylogenetic distribution of CHS-, CURS- and GS-associated metabolic activities across key nodal species spanning land plant evolutionary history. For each species, a filled square indicates that both a corresponding candidate gene was identified in the genome and the associated metabolic product was detected in available metabolomic data; open squares indicate that these combined criteria were not met. Green, yellow and red squares denote CHS-, CURS- and GS-associated activities, respectively. Selected genera within *Zingiberaceae* are highlighted. Triangles represent collapsed clades. **c)** Relative abundances of phenylpropanoid precursors, curcumin, 1-dehydrogingerdiones and gingerols in the rhizome, stem and leaf of ginger. Colour intensity represents the normalized metabolite peak area.4CL, 4-coumarate-CoA ligase; C3H, p-coumarate 3-hydroxylase; C3OMT, caffeic acid 3-O-methyltransferase.

Generally, metabolic innovation in plants often entails both the emergence of enzymes with new catalytic activities and the cell-type-specific coordination of specialized biosynthesis with precursor supply^18–20^. Ginger rhizomes contain specialized secretory cells, known as oil cells, in which pungent phenolic compounds preferentially accumulate in the cortex and pith^21–22^. This localized distribution suggests that gingerol biosynthesis may be concentrated within a small population of metabolically specialized cells rather than occurring uniformly across the rhizome. However, previous transcriptomic studies have largely relied on bulk tissues, in which signals from oil cells are averaged with those from surrounding cell types, potentially obscuring pathway genes expressed in these low-abundance cells^12–14^. Single-cell transcriptomic analysis therefore provides a means to resolve the oil-cell-associated metabolic program and to prioritize candidate enzymes from the expanded PKS and reductase families^23–26^.

Translating the logic of “micro-evolution” (gene sequence diversification and gene regulatory differentiation) leading to “macro-evolution” (metabolic lineage reconstruction) in natural evolution into rational design principles can provide evolutionary guidance and theoretical frameworks for synthetic biology research of natural products. Here, we integrated single-cell transcriptomics, functional screening, structural biology and comparative genomics to resolve the complete gingerol biosynthesis pathway and its evolutionary origin. Cell-resolved profiling identified the missing scaffold-forming and reductive enzymes in specialized oil-cells, whereas structural and mutational analyses revealed how limited active-site remodeling of a CURS-like type III PKS generated gingerol synthase activity. Translating these insights into yeast enabled the first de novo microbial production of 6-gingerol from glucose. Together, these findings establish the molecular basis of gingerol biosynthesis, demonstrate how cell-resolved expression information can guide the discovery and heterologous reconstruction of complex plant metabolic pathways, and reveal how limited remodeling of a type III PKS can expand its accessible chemical space. The resulting yeast platform provides a foundation to produce gingerols and structurally diversified analogues, while offering a broader framework for understanding lineage-specific metabolic innovation in plants.

## Results

### Determination of the catalytic order in the gingerol biosynthetic pathway

To determine the catalytic order in the gingerol biosynthetic pathway, we used metabolomic profiles of *Z. officinale* to constrain the synthetic network derived from retrosynthetic analysis (Supplementary Fig. 1a and b), thereby identifying the most likely catalytic route. We profiled pathway-related metabolites in the rhizome, stem and leaf. Among the predicted intermediates, high-abundance signals corresponding to 1-dehydrogingerdiones and the respective gingerols were detected, whereas all other intermediates in the predicted network were undetectable (Fig. 1c, Supplementary Fig. 1c). Thus, the gingerol biosynthetic pathway was established as follows: condensation of a feruloyl-CoA-derived β-keto intermediate with a fatty acyl-CoA assembles the 1-dehydrogingerdione scaffold, which is subsequently processed through ordered reduction to ultimately generate gingerols. This experimentally validated framework directed our search for the enzymes responsible for the unresolved scaffold-forming and reductive steps.

### Single-cell transcriptomics reveal a coordinated gingerol biosynthetic program

To identify the missing type III PKS and reductase in gingerol biosynthesis, we conducted genome-wide mining against the published *Z. officinale* genomes^13^ and identified 44 putative type III PKSs and 181 candidate reductases. Initially, bulk RNA sequencing of rhizomes, stems and leaves from one- and three-month-old ginger was processed to prioritize these candidates. However, differential-expression and co-expression analyses did not resolve a coherent candidate set (Supplementary Fig. 2a-c), indicating that gingerol biosynthesis might be restricted to a relatively rare cell population. We therefore performed high-throughput single-cell RNA sequencing on one-month-old ginger rhizome tissues, generating transcriptomes from 31,891 individual cells. Principal-component analysis (PCA) followed by UMAP embedding resolved 21 transcriptionally distinct clusters (Fig. 2a). We assigned cell identities—including epidermal, cortical, endodermal, pericycle, stele, bundle-sheath, vascular-bundle and oil cells by integrating orthologues of established marker genes from model plant with cluster-enriched genes and Gene Ontology enrichment analyses (Supplementary Fig. 3), and mapped their approximate spatial positions onto the rhizome anatomy.

**Fig. 2.**
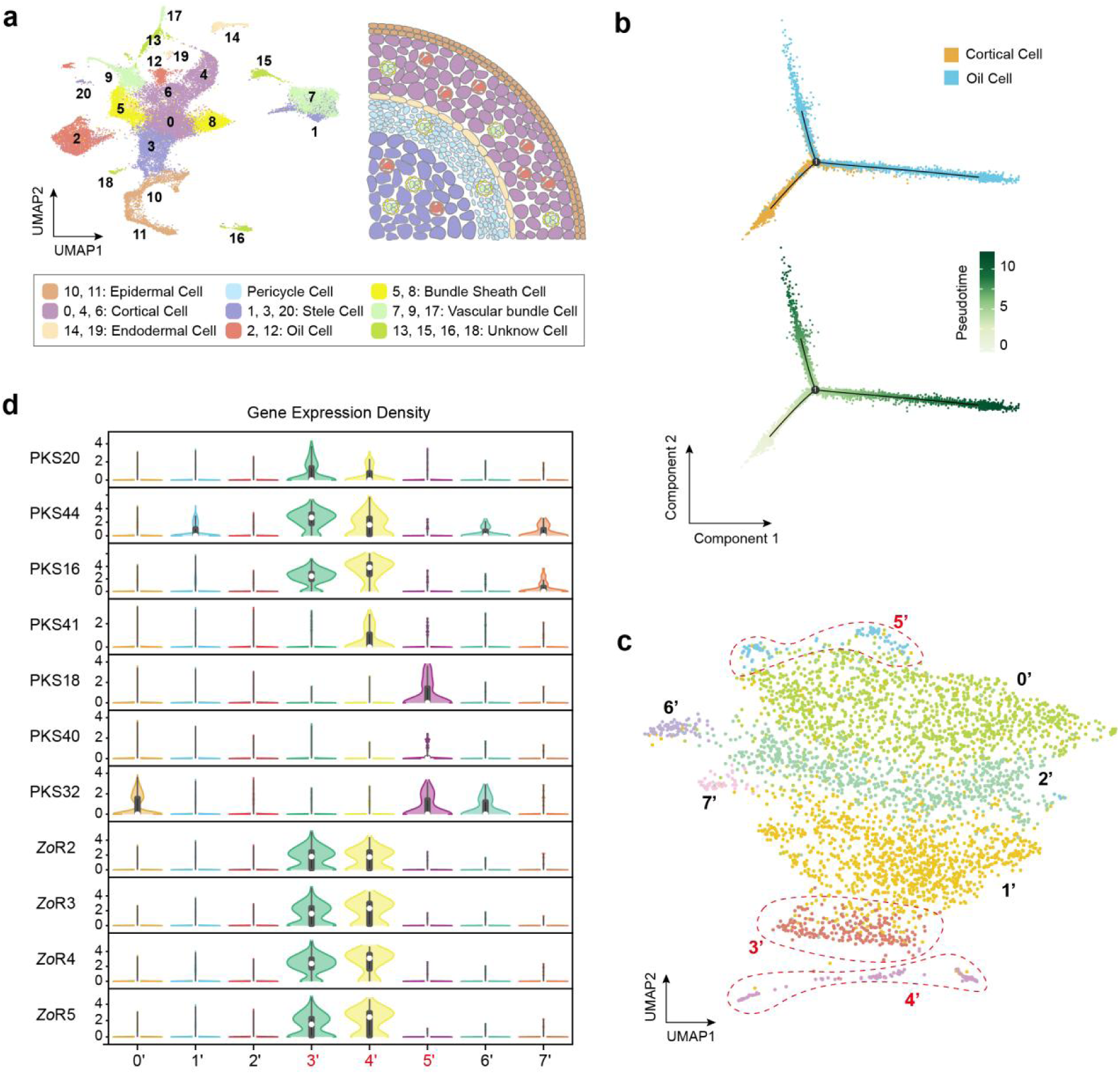
Single-cell transcriptomics identifies a metabolically specialized oil-cell population and candidate enzymes for gingerol biosynthesis. **a)** Uniform manifold approximation and projection (UMAP) of single cells isolated from *Z. officinale* rhizomes, resolving 21 transcriptionally distinct clusters. Cell identities were assigned on the basis of cluster-enriched marker genes and Gene Ontology enrichment analyses and are indicated by colour. The schematic illustrates the inferred spatial distribution of the annotated cell types within the rhizome tissue. **b)** Pseudotime analysis of cortical cells and oil cells using Monocle 2. The upper trajectory shows the distribution of cortical and oil cells along the inferred developmental branches. The lower trajectory is coloured by pseudotime, from early (light green) to late (dark green) states. Component 1 and Component 2 represent the reduced-dimensional coordinates used to reconstruct the cellular trajectory. **c)** UMAP representation of oil cells after subclustering into eight transcriptionally distinct subpopulations (0’-7’). Dashed outlines highlight subclusters enriched at the terminal regions of the oil-cell trajectory. **d)** Violin plots showing the expression distributions of candidate type III polyketide synthase genes and reductase genes across the eight oil-cell subclusters. Central dots and bars indicate the median and interquartile range, respectively.

Notably, transcriptomes of clusters 2 and 12 were distinguished by the enrichment of genes associated with polyketide and fatty acid metabolism (Supplementary Fig. 4a), which were expected to generate the aromatic and acyl precursors of gingerols. These clusters were therefore annotated as oil cells. This assignment was also consistent with previous histochemical studies that localized ginger pungent phenolics to specialized secretory cells distributed throughout the rhizome cortex and pith^21–22^. Therefore, 12 PKS genes and 6 reductase genes being upregulated in these clusters were considered candidates for gingerol biosynthesis (Supplementary Fig. 4b).

Pairwise pseudotime analysis revealed that only cortical cells formed a coherent continuous trajectory with oil cells (Fig. 2b), which was consistent with the fact that there was a close developmental relationship between the two cell lines. Cortical cells occupied earlier pseudotime positions, whereas oil cells segregated into two terminal branches at later positions, indicating that oil cells comprise at least two transcriptionally distinct terminal states rather than a uniform population. We therefore reclustered the oil cell population and resolved it into eight subclusters, designated 0’-7’ (Fig. 2c). Seven PKS genes, including PKS20, PKS44, PKS16, PKS41, PKS18, PKS40 and PKS32, exhibited subcluster-selective expression, so did four reductase candidates, including *Zo*R2, *Zo*R3, *Zo*R4 and *Zo*R5 (Fig. 2d). Notably, many of these candidates were preferentially expressed in subclusters 3’, 4’ and 5’. Meanwhile, genes encoding key enzymes of the phenylpropanoid pathway-including PAL, C4H, 4CL, C3H, CSE, COMT and CCOMT-and fatty-acyl substrate metabolism, including 3-ketoacyl-CoA thiolase, acyl-CoA oxidase and long-chain acyl-CoA synthetase, were specifically enriched in these subclusters (Supplementary Fig. 4c). This coordinated expression suggests that gingerol-forming enzymes are coupled to both aromatic and fatty-acyl precursor supply within metabolically active oil-cells, providing a credible candidate set for functional characterization.

### Identification of the core enzymes in the gingerol biosynthesis pathway

Gingerol scaffold assembly requires two consecutive type III PKS reactions, so we screened the single-cell-nominated candidates separately for DCS and GS activity. To identify GS, each of the seven PKS candidates was introduced into *S. cerevisiae* chassis expressing *At*4CL1^27^, *Cs*AAE1^28^ and *Cl*DCS^10^, which supplied feruloyl-β-keto acid and hexanoyl-CoA from exogenously added ferulic acid and hexanoic acid, respectively (Fig. 3a). Strains expressing PKS16, PKS40 and PKS44, respectively, predominantly produced curcumin, indicating that these PKSs possess CURS activity. None of the engineered strains accumulated the predicted GS product 1-dehydro-6-gingerdione, but either PKS18 or PKS20 accumulated certain amounts of curcumin and substantial amounts of 6-gingerol, indicating that PKS18 and PKS20 possess both CURS activities to form curcumin and the GS function to form the gingerol scaffold being subsequently reduced by endogenous yeast enzymes.

**Fig. 3.**
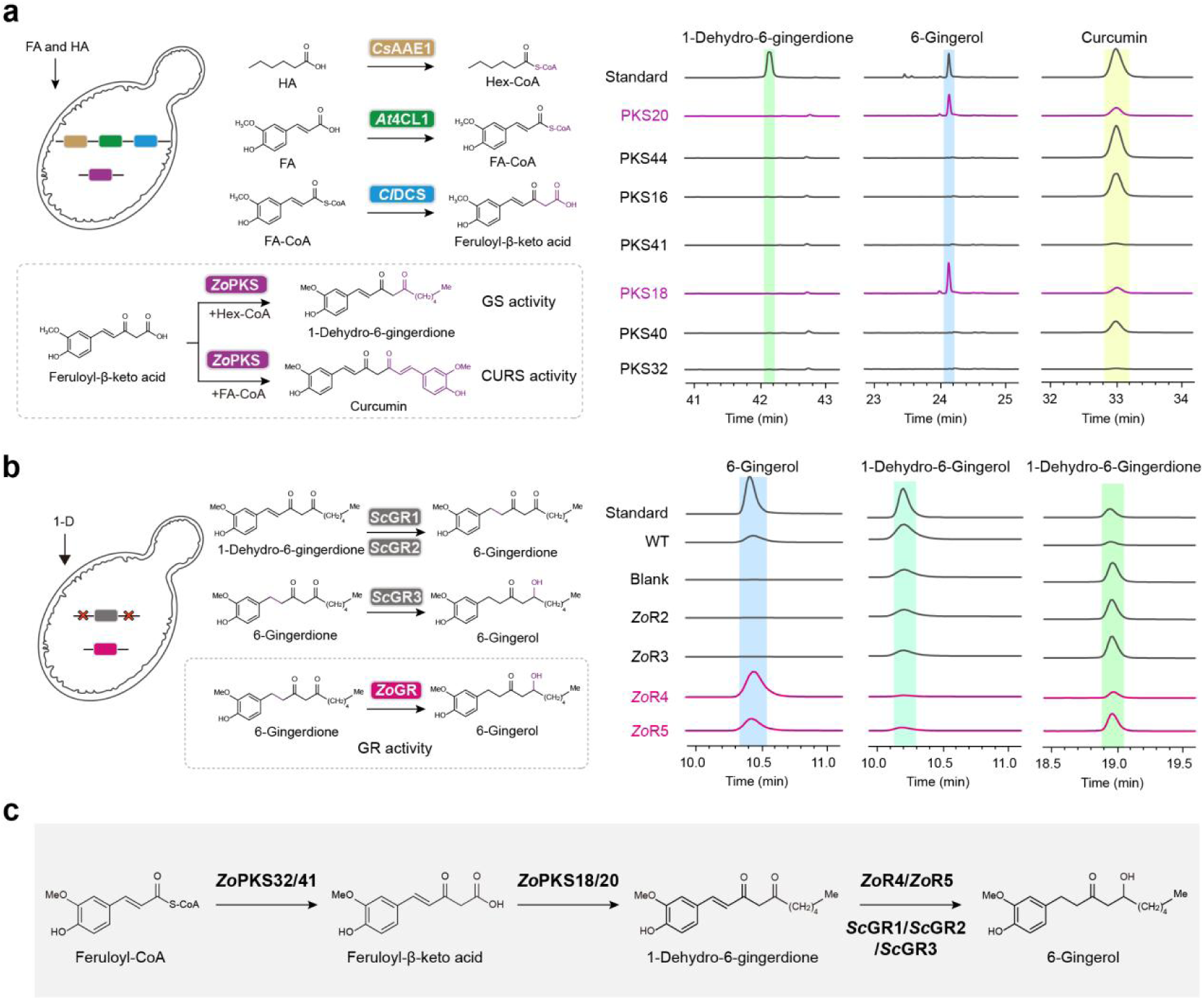
Functional screening identifies GS and GR activities that complete gingerol biosynthesis. **a)** Yeast-based screening of candidate type III PKS for GS and CURS activities. A screening chassis expressing *Cs*AAE1, *At*4CL and *Cl*DCS was constructed to generate hexanoyl-CoA and feruloyl-β-keto acid from exogenously supplied hexanoic acid (HA; 500 mM) and ferulic acid (FA; 75 mg l⁻¹). Candidate *Z. officinale* PKSs were individually expressed and assessed for their ability to condense feruloyl-β-keto acid with hexanoyl-CoA to form 1-dehydro-6-gingerdione, indicative of GS activity, or with feruloyl-CoA to form curcumin, indicative of CURS activity. Representative extracted-ion chromatograms show the production of 1-dehydro-6-gingerdione, 6-gingerol and curcumin by the engineered strains. Purple traces denote positive candidates, and dashed vertical lines indicate the retention times of authentic standards. **b)** Yeast-based screening of candidate *Zo*GRs. Yeast endogenous GRs reduce 1-dehydro-6-gingerdione to 6-gingerol, thereby confounding the identification of ginger GR activity. Three *Sc*GRs was therefore deleted to generate a screening chassis KO3 that accumulates 6-gingerdione. Candidate reductases were individually expressed and evaluated for conversion of 6-gingerdione into 6-gingerol. Representative extracted-ion chromatograms show 6-gingerol, 1-dehydro-6-gingerol and 1-dehydro-6-gingerdione accumulation in strains expressing the indicated candidates. WT, wild-type yeast; blank, empty-vector control in the KO3 chassis. Magenta traces indicate candidates exhibiting GR activity; dashed vertical lines denote the retention times of authentic standards. **c)** The core enzymatic pathway from feruloyl-CoA to 6-gingerol. 4CL, 4-coumarate-CoA ligase; AAE, acyl-activating enzyme; DCS, diketide-CoA synthase; FA-CoA, feruloyl-CoA; Hex-CoA, hexanoyl-CoA.

We then examined the PKS candidates catalyzed the preceding DCS reaction using *S. cerevisiae* chassis expressing *Cl*CURS^10^, and found that PKS32 and PKS41 had DCS activity (Supplementary Fig. 5a).

To identify yeast endogenous reductases capable of converting 1-dehydro-6-gingerdione into 6-gingerol, 1-dehydro-6-gingerdione was fed to the original yeast strain. The content of 1-dehydro-6-gingerdione progressively decreased, accompanied by the accumulation of 6-gingerdione initially and 6-gingerol subsequently (Supplementary Fig. 6a). Yeast therefore sequentially reduces the conjugated C=C bond of 1-dehydro-6-gingerdione and the carbonyl group of the resulting 6-gingerdione. This reaction order matches the pathway inferred from ginger tissue metabolite profiles (Fig. 1c), independently supporting 1-dehydrogingerdiones and gingerdiones as successive intermediates. By mining the yeast genome for putative oxidoreductases and constructing the knockout strains, we found that the deletion of either YDL114W (*Sc*GR1) or YMR152W (*Sc*GR2) increased the accumulation of 1-dehydro-6-gingerdione, and the deletion of YML131W (*Sc*GR3) caused 6-gingerdione accumulation without 6-gingerol formation (Supplementary Fig. 6b).

We therefore used the triple knockout *S. cerevisiae* strain KO3 (ΔScGR1 ΔScGR2 ΔScGR3) to screen for the ginger reductase candidates in the following study. The expression of either *Zo*R4 or *Zo*R5 in strain KO3 restored 6-gingerol production (Fig. 3b) and quantitative assays reflected that the 6-gingerdione converting efficiencies of *Zo*R4 and *Zo*R5 was higher than the endogenous yeast reductase (Supplementary Fig. 6c). Based on the above, we identified PKS32 and PKS41 as DCSs, PKS18 and PKS20 as GSs, and *Zo*R4 and *Zo*R5 as the terminal gingerol reductases, completing the core enzymatic pathway from feruloyl-CoA to gingerols (Fig. 3c).

### Structural basis of gingerol scaffold formation by GS

PKS18 and PKS20, the two enzymes with GS activity, share nearly identical amino acid sequences, differing at only a few residues outside the catalytic center (Supplementary Fig. 7a). We designated the more active PKS18 as *Zo*GS1 and selected it for structural and mechanistic characterization. X-ray crystallography resolved the apo structure of *Zo*GS1 at 2.5 Å resolution, revealing the canonical homodimeric architecture of type III polyketide synthases (Fig. 4a). The two protomers superimpose with an RMSD of 0.237 Å, indicating a near-symmetrical C2-related dimer (Supplementary Fig. 7b). Each protomer contain an independently accessible active-site cavity harbouring the conserved Cys169-His308-Asn341 catalytic triad characteristic of type III PKSs. The Met142 protrudes from the monomer surface across the dimer interface into a complementary hole in the adjoining monomer, forming part of the cyclization pocket. *Zo*GS1 shares the conserved dimeric fold and Cys-His-Asn catalytic machinery with other CHS-like enzymes, but local variations in active-site architecture determine substrate and product specificity. Residues 257–262 and 268–272 form a flexible region that may increase active-site plasticity and facilitate the accommodation of chemically distinct substrates.

**Fig. 4.**
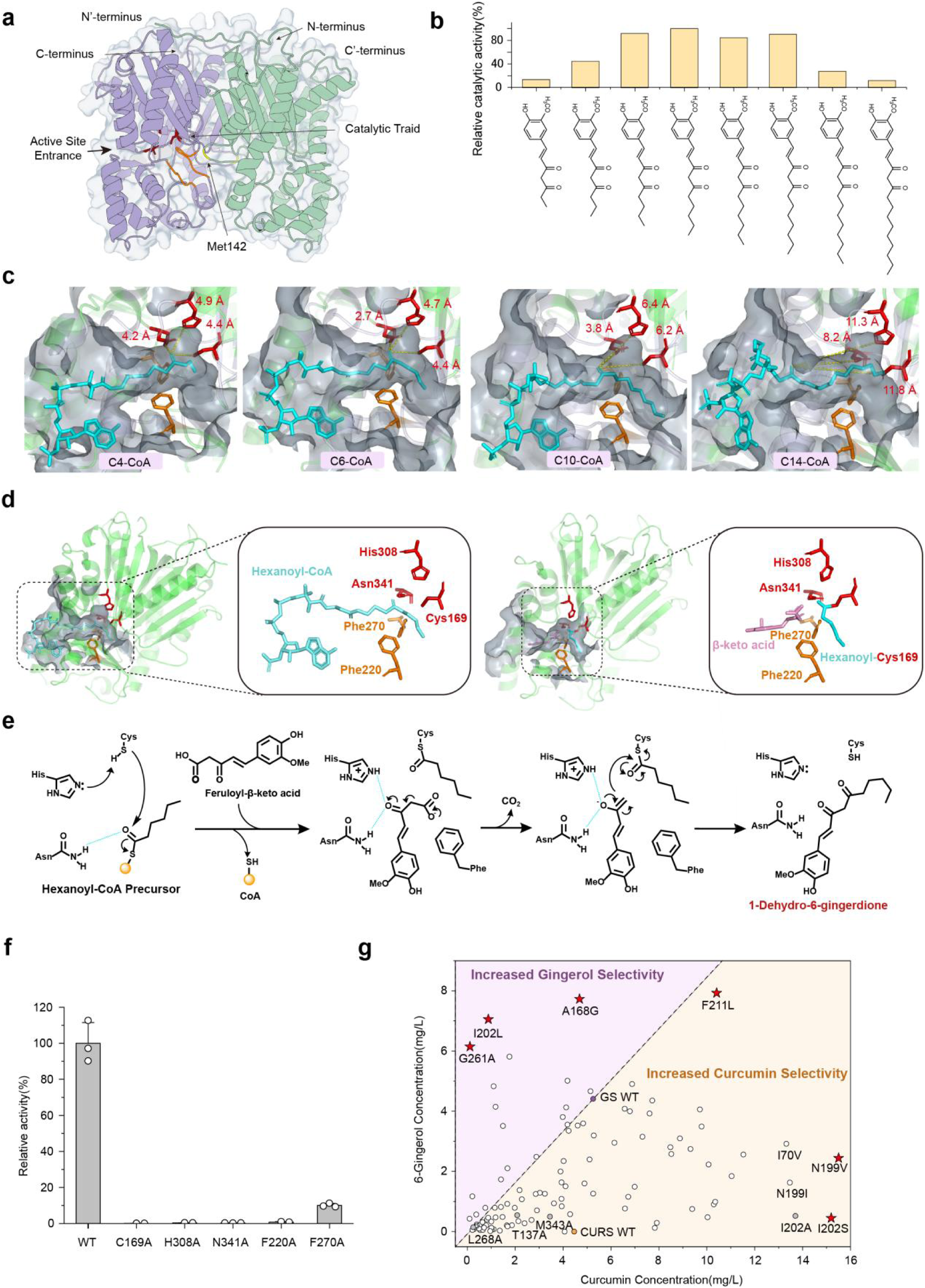
Structural and mutational basis of gingerol scaffold formation by GS. **a)** Crystal structure of *Zo*GS1, shown as a homodimer with the two subunits coloured purple and green. The active-site entrance, catalytic triad and Met142 are indicated. The disordered loop formed by residues 257–262 and 268–272 in the catalytic center is colored orange. The molecular surface is shown in transparent representation. **b)** Substrate profile of *Zo*GS1 towards acyl-CoA starter substrates of different chain lengths. Relative catalytic activities were determined from formation of the corresponding gingerol skeleton products and normalized to the activity towards the preferred substrate. **c)** Representative docking poses of acyl-CoA substrates bearing C4, C6, C10 and C14 acyl chains within the *Zo*GS1 active-site cavity. **d)** Molecular models of the two substrates involved in gingerol scaffold assembly following molecular docking and dynamics refinement. Left, the catalytic active center bound by the starting substrate hexanoyl-CoA. Right, binding of feruloyl-β-keto acid adjacent to the acylated catalytic cysteine. Insets show the spatial relationships of the substrates with Cys169, His308, Asn341 and Phe220. **e)** Proposed catalytic mechanism of PKS18. Cys169 is first acylated by hexanoyl-CoA, after which feruloyl-β-keto acid enters the active site and undergoes condensation with the enzyme-bound acyl intermediate. Decarboxylation and product release generate 1-dehydro-6-gingerdione. His308 and Asn341 form the conserved catalytic network, whereas surrounding pocket residues position the two substrates for productive coupling. **f)** Relative catalytic activities of *Zo*GS1 variants carrying substitutions at residues implicated in catalysis or substrate positioning. Activity of wild-type *Zo*GS1 was set to 100%. Three biological replicates were performed and data are presented as mean values ± SD. Individual points represent independent measurements. **g)** Characterization of *Zo*GS1 variants generated by alanine scanning and saturation mutagenesis of active-site residues. Each point represents one catalytically active variant and is positioned according to its activity of 6-gingerol and curcumin. The dashed diagonal separates variants with increased gingerol selectivity from those with increased curcumin selectivity. Wild-type GS and CURS are indicated, and selected variants with markedly altered activity or product preference are highlighted by red stars. Variants with no detectable activity are not shown. WT, wild type. Three biological replicates were performed and data are presented as mean values.

Substrate profiling shows *Zo*GS1 accepted several phenylpropanoid-derived substrates, producing gingerol-like products from p-coumarate-, dihydroferulate- and hydroxyphenylpropionate-related precursors (Supplementary Fig. 7c). It also accepts linear fatty acyl-CoAs of varying chain lengths, generating the corresponding 1-dehydrogingerdione homologues (Fig. 4b). By contrast, branched or otherwise modified acyl-CoAs showed little or no conversion. This broad tolerance towards linear acyl chains explains the homologous series of 4-, 6-, 8-, 10- and 12-gingerols in ginger tissues (Fig. 1c).

To understand how *Zo*GS1 accommodates different chain lengths, we docked C4, C6, C10 and C14 acyl-CoAs into the crystal structure and refined the complexes by molecular dynamics simulations (Fig. 4c). The active site contains an elongated hydrophobic channel (∼10 Å) extending beyond the catalytic cysteine. Linear acyl chains could occupy this channel while maintaining the thioester carbonyl near Cys169. For shorter substrates, CoA anchoring at the active-site entrance restricts acyl-chain flexibility, maintaining the thioester carbonyl within 5 Å of the catalytic triad in a correct competent orientation. Intermediate-chain substrates filled the hydrophobic channel more completely and maintained optimal positioning relative to the catalytic triad, consistent with their higher activities. In contrast, the C14 substrate exceeded the effective capacity of the channel, displacing its thioester carbonyl from the catalytic centre and providing a structural explanation for the diminished activity towards very long-chain substrates.

To model the *Zo*GS1-mediated scaffold formation steps, we first docked hexanoyl-CoA into *Zo*GS1 and generated the Cys169-bound hexanoyl intermediate, which we subsequently used to dock and refine feruloyl-β-keto acid by molecular dynamics (Fig. 4d). In the resulting model, the hexanoyl chain occupied the hydrophobic channel, and feruloyl-β-keto acid sits adjacent to the enzyme-bound thioester and the Cys169-His308-Asn341 catalytic triad. Phe220 and Phe270 act as gatekeepers that maintain active-centre hydrophobicity and orient the two substrates for C-C bond formation. Based on these models and the established mechanisms of CHS and CURS, we propose that *Zo*GS1 initiates catalysis through His308-assisted activation of Cys169, followed by nucleophilic attack on hexanoyl-CoA and transfer of the hexanoyl group to the catalytic cysteine with CoA release (Fig. 4e). After binding as the extender substrate, feruloyl-β-keto acid is decarboxylated to a nucleophilic intermediate, which condenses with the fatty acyl moiety to form 1-dehydro-6-gingerdione and restores the catalytic cysteine. This mechanism resembles the conserved starter-loading chemistry of CHS and the β-keto acid intermediate used by CURS, but *Zo*GS1 replaces the aromatic starter with a linear fatty acyl-CoA^10–11,29^.

Mutational analysis supported the proposed catalytic model. Substitution of Cys169, His308 or Asn341 nearly abolished *Zo*GS1 activity, confirming the essential roles of the conserved catalytic triad (Fig. 4f). Mutation of Phe220 also caused a pronounced loss of activity, consistent with its proposed role in organizing the substrates within the active-site cavity. Substitution of Phe270 produced a less severe but substantial reduction in activity (∼89.9% decrease), indicating that residues surrounding the catalytic center contribute to productive substrate positioning rather than directly mediating bond formation.These results support a mechanism in which the conserved type III PKS catalytic machinery combines with a distinct hydrophobic cavity to couple a fatty acyl-CoA starter with a phenylpropanoid-derived β-keto acid.

Finally, we investigated whether remodeling the *Zo*GS1 active-site pocket could alter its activity and product preference. Alanine scanning followed by saturation mutagenesis generated a broad spectrum of catalytically active variants, ranging from enzymes strongly biased towards gingerol formation to variants favouring curcumin production (Fig. 4g). The wild-type enzyme displayed bifunctional activity, producing both gingerol and curcumin. By contrast, G261A and I202L retained high gingerol-forming activity while producing little curcumin, increasing gingerol selectivity by 376.7-fold and 10.3-fold, respectively, relative to wild-type *Zo*GS1. F221L showed the highest gingerol-forming activity, reaching 1.8-fold that of the wild type. Conversely, I202S displayed curcumin-forming activity 3.4-fold higher than that of the characterized turmeric CURS used as the reference. Molecular dynamics simulations of G261A and I202L indicated the impact of pocket remodeling on catalytic selectivity (Supplementary Fig. 7e). Thus, limited remodeling of the *Zo*GS1 pocket is sufficient to separate its latent GS and CURS activities, identifying specific residues that govern the balance between two alternative type III PKS reaction outcomes.

### Gene duplication and catalytic remodeling create gingerol biosynthesis

Beside with the seven PKSs mentioned above, residual ginger PKSs prioritized by phylogenetic placement and single-cell expression patterns were functionally characterized in the study (Supplementary Fig. 4b). PKS1, PKS22 and PKS26 produced curcumin, being consistent with CURS activity, whereas PKS27, PKS42 and PKS43 supported feruloyl-β-keto intermediate formation, being consistent with DCS activity. These findings indicate that multiple functionally related PKS copies exist in ginger. Since these PKSs represent allelic counterparts on the diploid ginger genome, we analyzed their genomic organization within a single haplotype to avoid conflating allelic redundancy with gene duplication (Fig. 5a). Strikingly, the functionally characterized genes were organized into spatially associated GS-DCS and CURS-DCS pairs within the ginger genome.

**Fig. 5.**
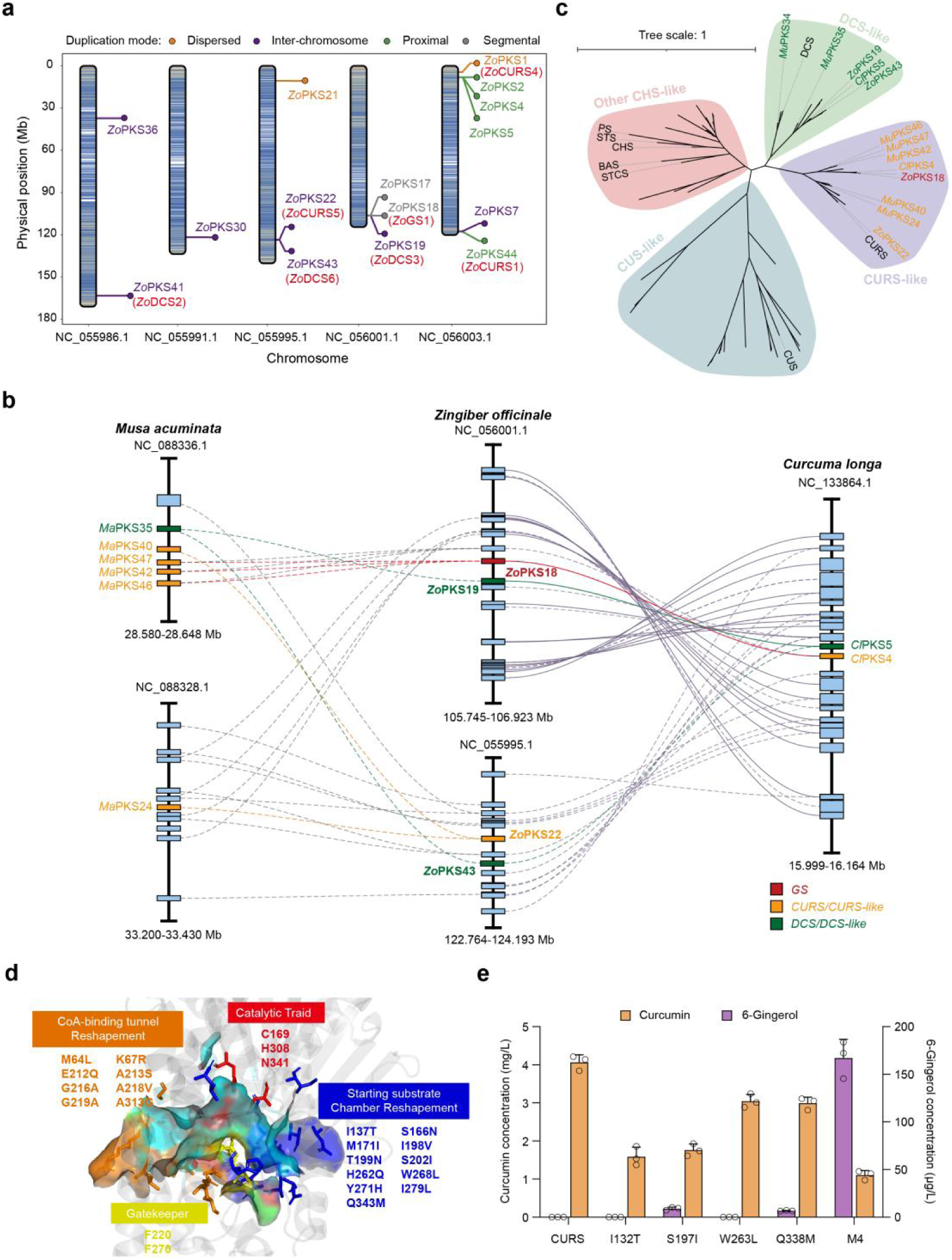
Gene duplication and catalytic remodeling underlie the emergence of gingerol biosynthesis. **a)** Chromosomal distribution and inferred duplication modes of type III PKS genes in the haplotype-resolved ginger genome. PKS loci are positioned according to their physical coordinates and coloured by duplication category. **b)** Local syntenic relationships among PKS-containing chromosomal regions in banana, ginger and turmeric. Rectangles denote genes, and connecting lines indicate homologous gene pairs between species. Experimentally validated GS, CURS and DCS genes from ginger and turmeric are highlighted in red, orange and green, respectively. Banana PKSs are colored according to their predicted DCS- or CURS-like annotations and have not been functionally characterized. **c)** Maximum-likelihood phylogeny of type III PKSs identified from *Z. officinale*, *C. longa* and *M. acuminata*, together with functionally characterized members of the chalcone synthase superfamily. Major clades corresponding to DCS-like, CURS-like, CUS-like and other CHS-like enzymes are shaded. Experimentally characterized ginger and turmeric PKSs are labelled according to their validated activities, whereas banana PKSs are annotated as putative DCS- or CURS-like proteins on the basis of sequence similarity, phylogenetic placement and available protein annotations. Branch lengths represent amino-acid substitutions per site. **d)** structural mapping of amino-acid differences between *Zo*GS1 and *Cl*CURS onto the *Zo*GS1 active-site cavity. **e)** Product profiles of *Cl*CURS variants carrying GS-associated amino-acid substitutions. Curcumin and 6-gingerol production by wild-type CURS, the single-site variants I132T, S197I, W263L and Q338M, and the combined four-site mutant M4 were quantified in yeast. Three biological replicates were performed and data are presented as mean values ± SD.

Local synteny analysis revealed CURS–DCS-like gene combinations in banana (*Musa acuminata*) and turmeric (*Curcuma longa*) corresponding to those in ginger (*Zingiber officinale*), with extensive collinearity among PKS-containing regions across all three species (Fig. 5b). The ginger GS-DCS region was syntenic with CURS-DCS-containing regions, suggesting that these functionally distinct modules originated from related ancestral genomic segments. Notably, the putative banana CURS- and DCS-like genes occurred within a local array of seven tandemly duplicated PKSs^30^ (Supplementary Fig. 8a). The banana proteins were assigned as CURS- or DCS-like based on annotation, sequence similarity and genomic context and were not biochemically characterized. Nevertheless, their organization is consistent with tandem expansion of a CHS-related locus followed by functional divergence of individual PKS copies.

To further evaluate this hypothesis, we reconstructed the phylogenetic relationships of all PKSs identified from banana, turmeric and ginger together with representative characterized type III PKSs from other species (Fig. 5c). DCS-like and CURS-like proteins formed two distinct phylogenetic groups, indicating that the complementary activities required for diarylheptanoid biosynthesis arose through divergence of separate PKS lineages. Ginger GSs clustered within the broader CURS-like group rather than with DCS-like enzymes. Combined with the conserved syntenic relationships between the GS-DCS and CURS-DCS regions, this clustering supports a evolutionary genetic hypothesis in which that an ancestral CHS-related tandem expansion produced a DCS-CURS-like module, followed by the further functional divergence of a CURS-like copy to generate GS activity in the ginger lineage.

We therefore investigated how a CURS-like enzyme acquires gingerol-forming activity. *Zo*GS1 retains the conserved catalytic triad of *Cl*CURS, but residues surrounding the CoA-binding tunnel, gatekeeper region and starter-substrate chamber differ between the two enzymes (Fig. 5d). By Combining this structural comparison with the *Zo*GS1 mutational landscape, we identified four positions-137, 202, 268 and 343 in *Zo*GS1-whose substitution substantially reduced gingerol formation (Fig. 4g). So, we designated these as candidate determinants of the functional distinction between *Zo*GS1 and CURS. We introduced the corresponding substitutions into *Cl*CURS, generating four single variants I132T, S197I, W263L and Q338M, and one four-site variant M4. S197I and Q338M individually conferred detectable GS activity on the CURS scaffold (Fig. 5e), demonstrating that single amino-acid changes could unlock latent GS activity in a CURS background. The M4 variant caused a more pronounced shift in product distribution, markedly increasing 6-gingerol formation while reducing curcumin production. These findings identify key amino-acid substitutions that drive the CURS-to-GS functional transition and demonstrate that remodeling of the substrate-binding pocket, rather than alteration of the conserved catalytic machinery, enabled the emergence of gingerol-forming activity.

Genomic and biochemical evidence convergently supports a stepwise model in which tandem duplication and divergence of CHS-related genes generated complementary DCS- and CURS-like functions in *Zingiberales*, and subsequent active-site remodelling of a CURS-like paralogue produced GS activity and established the lineage-specific GS-DCS module of gingerol biosynthesis.

### Synthetic reconstruction establishes de novo 6-gingerol biosynthesis in yeast

To reconstruct the gingerol biosynthesis pathway in yeast and achieve efficient cell factory synthesis, we first developed a series of strains with progressively enhanced ferulic acid production (Fig. 6a and b). Building on reported strategies to enhance the biosynthesis of ferulic acid^31–33^, we sequentially introduced TAL- and PAL-dependent phenylpropanoid modules, deleted PAD1 to limit endogenous phenylacrylic-acid decarboxylation, expressed feedback-insensitive aromatic-pathway enzymes and installed a phosphoketolase-based carbon-rewiring module. These modifications generated strains FA01-FA05 with progressively increased production of p-coumaric acid, caffeic acid and ferulic acid (Fig. 6b, Supplementary Fig. 9a).

**Fig. 6.**
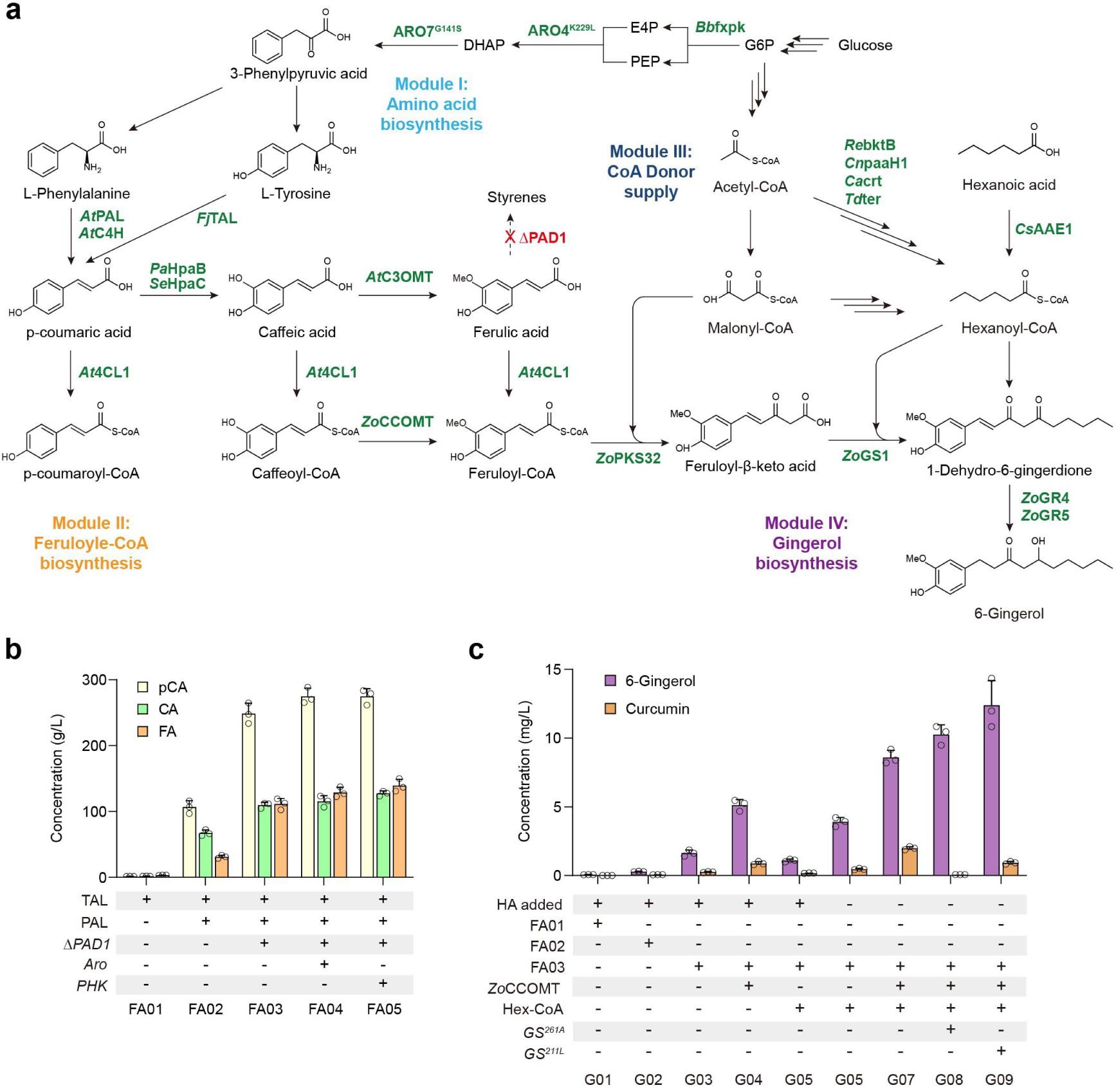
Gingerol biosynthesis module assembly and de novo production in yeast. **a)** Schematic of the engineered gingerol biosynthetic pathway in yeast. The pathway was organized into four modules: enhancement of aromatic amino-acid biosynthesis from glucose (module I); conversion of phenylalanine or tyrosine into feruloyl-CoA, accompanied by deletion of PAD1 to suppress byproduct formation (module II); reinforcement of hexanoyl-CoA supply (module III); and assembly of the gingerol pathway using ginger DCS (ZoDCS1), GS (ZoGS1) and endogenous ScGRs (module IV). Green labels indicate introduced or engineered enzymes, red crosses denote gene deletions, and dashed boundaries define the four engineering modules. **b)** Stepwise enhancement of phenylpropanoid precursor production in strains FA01–FA05. Titres of p-coumaric acid (pCA), caffeic acid (CA) and ferulic acid (FA) were quantified after sequential introduction of phenylalanine ammonia-lyase based pathway (PAL), tyrosine ammonia-lyase based pathway (TAL), deletion of PAD1, reinforcement of aromatic amino-acid biosynthesis (Aro), and phosphoketolase-based carbon redistribution (PHK). The matrix below the graph indicates the engineering components present in each strain. Three biological replicates were performed and data are presented as mean values ± SD. **c)** Stepwise optimization of gingerol biosynthesis in strains G01–G08. Production of 6-gingerol and the competing product curcumin was evaluated following incorporation of the FA01–FA03 precursor-producing modules, hexanoic acid feeding (HA), reinforcement of feruloyl-CoA (*Zo*CCOMT) and hexanoyl-CoA (Hex-CoA) supply, and introduction of the high-performance GS18 variants G261A and I211L. The plus–minus matrix indicates the engineering features present in each strain. Three biological replicates were performed and data are presented as mean values ± SD. G6P, glucose 6-phosphate; E4P, erythrose 4-phosphate; PEP, phosphoenolpyruvate; DHAP, dihydroxyacetone phosphate.

We next introduced the core gingerol pathway (*Zo*DCS1 and *Zo*GS1) into strains with low, intermediate and high ferulic-acid production. In the presence of 500 μM exogenous hexanoic acid, all three chassis produced 6-gingerol, and titres increased with the capacity of the host to supply phenylpropanoid precursors (Fig. 6c). These results established the functional reconstruction of gingerol biosynthesis in yeast and indicated that precursor availability remained a major determinant of pathway output. We then introduced *Zo*CCOMT, a highly expressed candidate identified from the ginger single-cell transcriptome, to enhance the conversion of caffeic acid-derived intermediates towards feruloyl-CoA (Fig. 6a and Supplementary Fig. 4c). Incorporation of *Zo*CCOMT markedly increased 6-gingerol production by 214.7% (Fig. 6c).

Substrate-feeding experiments showed that increasing hexanoic-acid availability strongly stimulated gingerol formation, identifying hexanoyl-CoA supply as another bottleneck. We therefore introduced a heterologous reverse β-oxidation module (*Re*BktB, *Cn*PaaH1, *Ca*Crt and *Td*Ter) to generate hexanoyl-CoA from endogenous acetyl-CoA^34^ (Fig. 6a). This module enabled substantial 6-gingerol production in the absence of exogenous hexanoic acid, completing the de novo synthesis from glucose. Notably, additional hexanoic-acid supplementation decreased rather than increased gingerol titers in the engineered strain. This response aligns with excess hexanoate or hexanoyl-CoA imposing cellular stress or disrupting the balance between the two GS substrates.

Finally, we combined enhanced feruloyl-CoA and hexanoyl-CoA supply with *Zo*GS1 variants selected for improved activity or gingerol preference. Introduction of the high-performance GS variants further increased carbon flux towards 6-gingerol while suppressing formation of the competing product curcumin (Fig. 6c). The resulting strain produced 12.4 mg/L 6-gingerol without precursor supplementation.

## Discussion

Here, we resolve the complete biosynthetic pathway of gingerols by identifying the consecutive DCS, GS and GR reactions responsible for scaffold assembly and reduction. Heterologous reconstruction further demonstrated that these enzymes, together with the required precursor-supply pathways, are sufficient to generate gingerol from glucose. To our knowledge, this represents the first de novo microbial biosynthesis of gingerol. Beyond validating the proposed pathway, the engineered yeast system provides a platform for producing gingerols and structurally diversified analogues with different fat chain lengths and branched chains, as demonstrated for other complex plant natural products reconstructed in microbial hosts^4,35–36^.

Single-cell transcriptomics was central to resolving this pathway. Whereas bulk-tissue analyses could not effectively distinguish functional enzymes from the expanded PKS and reductase families, cell-resolved expression identified oil-cell subpopulations in which phenylpropanoid metabolism, fatty acyl-CoA supply and gingerol-forming enzymes were coordinately enriched. This analysis guided the identification of DCS, GS and GR enzymes and also revealed the high-performance precursor-supply enzyme *Zo*CCOMT, which enhanced gingerol production in yeast. These findings highlight the value of single-cell approaches not only for locating specialized metabolic programmes and identifying pathway enzymes, but also for selecting efficient components for pathway reconstruction^24,37–38^. The coordinated activation of precursor-supply and product-forming reactions further suggests that the emergence of gingerol biosynthesis required regulatory integration of an entire metabolic programme within specialized cells. Whether these oil-cell states are governed by dedicated transcriptional or metabolic regulatory circuits that optimize pathway flux and product accumulation remains an important question for future investigation.

*Zo*GS1 exemplifies the pronounced catalytic plasticity of type III PKSs^39–42^. While retaining the conserved catalytic machinery of CHS-like enzymes, it organizes two chemically distinct building blocks—a fatty acyl-CoA and a phenylpropanoid-derived β-keto acid—into the gingerol scaffold. The crystal structure of *Zo*GS1 reveals the molecular features that govern substrate accommodation and positioning, providing a structural blueprint for engineering chain-length preference, catalytic activity and product selectivity. Its broad acceptance of linear acyl-CoAs also provides a biochemical explanation for the natural diversity of gingerol homologues. Notably, *Zo*GS1 generated odd-chain gingerol analogues that are not normally detected in ginger, suggesting that the predominance of even-chain gingerols in planta is determined at least partly by the composition of the intracellular acyl-CoA pool rather than by strict enzymatic exclusion. Thus, enzyme promiscuity defines the chemical space accessible to GS, whereas precursor availability determines which products are realized in the native plant. Whereas chemical synthesis enables precise access to individual gingerol congeners, biosynthesis offers a complementary, modular strategy in which precursor variation and structure-guided enzyme engineering can be combined to generate families of natural and non-natural analogues under biologically compatible conditions.

Our genomic, phylogenetic and biochemical analyses support a model in which gingerol biosynthesis arose through both catalytic and regulatory evolution. Duplication and divergence of CHS-related genes probably generated complementary DCS- and CURS-like functions, followed by active-site remodelling of a CURS-like paralogue to produce GS activity. The acquisition of gingerol-forming activity by engineered CURS variants demonstrates that limited amino-acid substitutions can redirect this conserved PKS scaffold towards a new product. However, the appearance of GS alone would not have been sufficient to establish gingerol biosynthesis; its integration with DCS, GR and the coordinated supply of feruloyl-CoA and fatty acyl-CoAs was also required. Translating these natural principles into yeast enabled the first de novo reconstruction of gingerol biosynthesis and demonstrated that cell-resolved expression information can guide the selection of enzymes for pathway improvement. More broadly, the combination of single-cell pathway discovery, evolutionary analysis and structure-guided enzyme engineering offers a route not only to reconstruct lineage-restricted plant metabolites, but also to extend their chemical space beyond that realized by the native producer.

## Methods

### Phylogenetic analysis

Phylogenetic analysis was performed using shared single-copy plastid protein-coding genes. Plastid CDSs were extracted from GenBank files using PhyloSuite^43^, and single-copy orthologous genes shared by all species and showing consistent annotations were identified using OrthoFinder^44^. Genes with missing sequences, pseudogenization, ambiguous boundaries, abnormal lengths or unresolved annotation inconsistencies were excluded. Each CDS was translated and aligned separately at the amino acid level using the L-INS-i algorithm in MAFFT, followed by inspection for truncated or duplicated sequences, extensive gaps, misalignment and unusually divergent sequences. Poorly aligned regions were removed using trimAl with the parameter -gt 0.5, and problematic alignments were re-examined or excluded. The trimmed alignments were concatenated into a partitioned amino acid supermatrix using AMAS. Maximum-likelihood phylogenetic inference was performed using IQ-TREE 2^45^ with the parameters -m MFP+MERGE -bb 1000 -alrt 1000, allowing automatic model selection and partition merging, with branch support assessed using 1,000 ultrafast bootstrap replicates and 1,000 SH-aLRT replicates.

### Metabolomic profiling of ginger

Dried rhizome, stem and leaf tissues of ginger were ground into fine powders under liquid nitrogen. Accurately weighed samples were extracted with ethyl acetate, concentrated to dryness, redissolved in methanol and filtered through a 0.22-μm membrane before analysis. Metabolite profiling was performed using an Ultimate 3000 UHPLC system coupled to a Q Exactive Plus Orbitrap high-resolution mass spectrometer equipped with a heated electrospray ionization source (Thermo Fisher Scientific, USA). Chromatographic separation was conducted on an ACQUITY UPLC HSS T3 C18 column (2.1 × 100 mm, 1.8 μm; Waters, USA) maintained at 35 °C, with 0.1% (v/v) formic acid in water and 0.1% (v/v) formic acid in acetonitrile as mobile phases A and B, respectively. The flow rate was 0.2 mL/min and the injection volume was 5 μL. The gradient was as follows: 0–10 min, 100% B; 10–20 min, 100–70% B; 20–25 min, 70–60% B; 25–30 min, 60–50% B; 30–40 min, 50–30% B; 40–45 min, 30–0% B; 45–60 min, 0% B; 60–60.1 min, 0–100% B; and 60.1–70 min, 100% B. Mass spectra were acquired in both positive- and negative-ion modes, with sheath and auxiliary gas flow rates of 40 and 15 arbitrary units, respectively. The capillary and auxiliary gas heater temperatures were set to 320 and 350 °C, respectively, and the spray voltages were +3.2 kV and −3.0 kV in positive-and negative-ion modes, respectively. Full-scan spectra were acquired over an m/z range of 100–1,500 at a resolution of 70,000, followed by data-dependent MS/MS acquisition using higher-energy collisional dissociation at a resolution of 17,500. Metabolites were annotated using Compound Discoverer 3.3 against the mzCloud and mzVault databases, with a precursor mass tolerance of ±5 ppm. Peak areas were normalized to sample weight and used for relative metabolite comparisons and heatmap visualization; no absolute quantification was performed.

### Retrobiosynthesis analysis

A bio-retrosynthesis workflow was used to predict potential biosynthesisc routes to 6-gingerol. Structural analogues of 6-gingerol were first identified among KEGG metabolites by molecular fingerprint similarity analysis using RDKit. Potential pathways connecting these analogues to 2,808 endogenous metabolites extracted from the Saccharomyces cerevisiae genome-scale metabolic model Yeast8^46^ were subsequently searched using the shortest_simple_paths algorithm implemented in NetworkX. RetroRules^47^, comprising 65,688 reaction rules with diameters of 4 and 6, was then applied to predict retrosynthesis transformations from 6-gingerol to the identified structural analogues. To constrain combinatorial expansion, similarity-guided Monte Carlo tree search was employed to identify plausible retrosynthesis routes. The resulting routes were further expanded through human-guided single-step retrosynthetic analysis and manually curated to construct the final retrosynthesis network.

### Bulk RNA sequencing of ginger organs

Rhizome, stem and leaf tissues were collected from one- and three-month-old ginger plants. Total RNA was extracted using TRIzol reagent, and 1 μg of RNA per sample was used for library construction. Poly(A)+ mRNA was enriched using oligo(dT) beads and fragmented using divalent cations at elevated temperature. First-and second-strand cDNA synthesis was performed using random primers, followed by end repair, dA tailing and adapter ligation. Adapter-ligated fragments were size-selected using DNA purification beads and amplified using P5 and P7 primers. Indexed libraries were pooled and sequenced on an Illumina HiSeq platform in paired-end 150-bp mode. After quality filtering, clean reads were aligned to the ginger reference genome using HISAT2 v2.2.1. Gene models were predicted using GeMoMa with genomic and transcriptomic evidence from closely related species, and annotation completeness was assessed using BUSCO. Transcript abundance was quantified using StringTie.

### Single-cell RNA sequencing of ginger rhizome

#### Zingiber officinale cv

Zhangliang was used as the experimental material. Rhizomes were collected from healthy, pest- and disease-free plants one month after planting. After the roots and buds were removed, the rhizomes were thoroughly rinsed with clean water and immediately used for nuclei isolation. Single-nucleus RNA sequencing was performed by Gene Denovo Biotechnology Co., Ltd. (Guangzhou, China) using the 10× Genomics platform. Crude nuclei were isolated and subjected to quality assessment, and qualified suspensions were adjusted to 1,000–2,000 nuclei μL⁻¹. Nuclei were mixed with barcoded gel beads and reverse-transcription reagents and encapsulated into Gel Beads-in-Emulsion droplets. Following reverse transcription and barcode incorporation, the emulsions were disrupted, and the resulting cDNA was amplified, pooled and enzymatically fragmented to approximately 200–300 bp. Sequencing libraries were prepared by end repair, A-tailing, ligation of P5 and P7 adapters, sample indexing and PCR amplification, and were sequenced on an Illumina platform in paired-end 150-bp mode.

Raw reads were processed using Cell Ranger and mapped to the ginger reference genome assembly GCA_018446385.1. Subsequent quality control and analysis were performed using Seurat. Doublets were identified and removed using DoubletFinder^48^. Nuclei with fewer than 270 or more than 7,200 detected genes, more than 11,000 unique molecular identifiers, or chloroplast-derived transcript proportions outside the range of 0–0.31% were excluded. Gene-expression values were normalized using the LogNormalize method, and 2,000 highly variable genes were selected. Batch effects were corrected using Harmony^49^, followed by principal component analysis and Louvain clustering at a resolution of 0.5. Cell clusters were visualized using Uniform Manifold Approximation and Projection. Differentially expressed genes were identified using the Wilcoxon rank-sum test with thresholds of P < 0.01, |log₂ fold change| > 0.5 and min.pct > 0.25. Cell types were annotated according to cluster-specific marker genes and their homologues in *Arabidopsis thaliana*, *Oryza sativa*, *Zea mays* and *Glycine max*. Gene Ontology and KEGG enrichment analyses were performed using the OmicShare platform.

### Gene expression cassette and strain construction

Promoters, terminators, open reading frames and homologous arms were amplified by PCR using 2× Phanta Flash Master Mix (Vazyme, China). PCR products were separated by 1% agarose gel electrophoresis and purified using the GeneJET Gel Extraction Kit (Thermo Fisher Scientific). DNA fragments were assembled into expression cassettes by overlap-extension PCR or Gibson assembly. The resulting constructs, flanked by homologous arms, were introduced into *Saccharomyces cerevisiae* BY4742 cells by electroporation and integrated into the genome through homologous recombination. Transformants were selected on YPD agar plates at 30 ℃ for 2 days, and positive colonies were confirmed by diagnostic PCR.

### Strain fermentation

Single yeast colonies were inoculated in triplicate into 2 mL YPD medium and cultured overnight to saturation at 30 ℃ and 400 rpm. Subsequently, 100 μL of each culture was transferred into 4 mL YPD medium in 24-well deep-well plates sealed with breathable sealing films (Biotss). Cultures were incubated for 120 h at 30 ℃ and 400 rpm in a high-speed orbital shaker (ZSZY-88BH; Shanghai Zhichu Instrument Co., Ltd.). After 24 h, galactose was added to a final concentration of 20 g/L to induce gene expression, together with the indicated substrates when required.

### Analysis of metabolite production

Fermentation cultures were disrupted by bead beating and extracted with two volumes of ethyl acetate. The organic phase was collected, evaporated to dryness and reconstituted in methanol for LC–MS analysis. Metabolite production was quantified using a Shimadzu LCMS-8040 system equipped with an ACQUITY Premier HSS T3 column (2.1 × 100 mm, 1.8 μm). The column temperature was maintained at 40 °C. Mobile phases A and B were 0.1% (v/v) formic acid in water and acetonitrile, respectively. The gradient was as follows: 0 min, 20% B; 0–5 min, 20–30% B; 5–7 min, 30–42% B; 7–12 min, 42–45% B; 12–14 min, 45–50% B; 14–16 min, 50–60% B; 16–19 min, 60–70% B; 19–23 min, 70–100% B; and 23–26 min, 100–20% B. The flow rate was 0.4 mL min⁻¹, and the injection volume was 3 μL.

### Protein expression, purification, and crystallization

The coding sequence of *Zo*GS1 was cloned into pET-28a with an N-terminal 6×His tag and a TEV protease cleavage site. Recombinant cells were cultured in kanamycin-supplemented TB medium at 37 °C and 220 rpm, and protein expression was induced with 0.6 mM IPTG at 16 °C for 36 h. Cells were harvested and disrupted by high-pressure homogenization in 50 mM Tris–HCl (pH 8.0), 300 mM NaCl, 20 mM imidazole, 10 mM β-mercaptoethanol and 10% glycerol. The clarified lysate was subjected to Ni-affinity chromatography, followed by TEV cleavage, reverse Ni-affinity chromatography, Q anion-exchange chromatography and size-exclusion chromatography. Purified GS18 was concentrated to approximately 10 mg mL⁻¹ in 20 mM Tris–HCl (pH 8.0), 150 mM NaCl and 2 mM TCEP.

Crystallization was performed at 4 °C using the sitting-drop vapour-diffusion method by mixing 100 nL protein solution with 100 nL reservoir solution and equilibrating the drops against 35 μL reservoir solution. Diffraction-quality apo *Zo*GS1 crystals were obtained under the ProPlex A10 condition. Crystals were cryoprotected in reservoir solution supplemented with 10% glycerol, and X-ray diffraction data were collected at beamline BL10U2 of the Shanghai Synchrotron Radiation Facility. Diffraction images were indexed and integrated using XDS and merged and scaled using AIMLESS. The structure was solved by molecular replacement using Phaser, manually rebuilt in Coot and refined using phenix.refine.

### Substrate profiling

Substrate specificity was evaluated *in vivo* using engineered *Saccharomyces cerevisiae* strains expressing *Zo*GS1 together with following heterologous CoA ligases: *At*4CL1 from *Arabidopsis thaliana*^27^, *Ec*FadD from *Escherichia coli*^50^, *Hl*CCL2/*Hl*CCL4 from *Humulus lupulus*^51^, and *Cs*AAE1 from *Cannabis sativa*^28^. These enzymes converted exogenously supplied phenylpropanoid-derived acids and straight- or branched-chain fatty acids into the corresponding CoA thioesters, thereby supplying alternative starter units for PKS catalysis. The test substrates were added to the induced yeast cultures at the indicated concentrations. Cultivation, metabolite extraction and LC–MS analysis were performed as described in **Media and strains** and **Analysis of metabolite production**.

### Covalent docking of the reaction intermediate

The experimentally determined apo *Zo*GS1 crystal structure was used for docking. Crystallographic waters and additives were removed, missing side chains were completed, and hydrogen atoms were added at pH 7.4. A hexanoyl-derived C6 acyl fragment was first docked into the apo active site using flexible-residue docking in smina, thereby defining its orientation relative to the catalytic residue Cys169.

A covalent acyl–enzyme intermediate representing the thioester state preceding Claisen condensation was then constructed from the selected pose. The thiol hydrogen of Cys169 was removed, the residue was assigned as CYX, and a covalent bond was introduced between Sγ of Cys169 and the acyl carbonyl carbon with an equilibrium length of 1.81 Å. The complex was energy-minimized in GROMACS^52^ using the CHARMM36^53^ force field, with acyl-fragment parameters generated using CGenFF. The covalent bond and associated angle and dihedral terms were incorporated into the topology. Steepest-descent minimization was performed first with the protein backbone restrained at 1,000 kJ mol⁻¹ nm⁻² and subsequently without restraints.

Feruloyl β-keto acid was docked into the minimized covalent intermediate using smina, with the covalently linked acyl fragment treated as part of the receptor. Cys169, His271, His308, Ser337, Glu338 and Asn341 were allowed to move, whereas the remaining receptor was held rigid. The docking box was centred at (2.56, 0.21, −6.91 Å) with dimensions of 25×30×20 Å. Docking was performed in 10 independent runs with an exhaustiveness of 16 and up to 20 poses per run using distinct random seeds and enhanced hydrogen-bond weighting. Productive poses were selected according to catalytic geometry rather than docking score alone, requiring orientation of the β-keto oxygen toward His308 and Asn341 and proximity of the ligand α-carbon to the thioester carbon.

### Structure prediction and molecular dynamics simulations

Three-dimensional enzyme–substrate complex models were generated using Boltz-2^54^. A contact restraint was imposed between the catalytic cysteine sulfur atom (Cys169 Sγ) and the substrate thioester acyl carbon to favour catalytically competent Michaelis complexes. Five models were generated per system and evaluated using the Boltz-2 confidence score, pTM, ipTM and complex pLDDT. The proportion of models with a Cys(Sγ)–acyl-C distance ≤4.0 Å and the minimum distance observed were recorded, and the highest-confidence model was selected for molecular dynamics simulation.

Each complex was placed in a rhombic dodecahedral box with a minimum solute-to-box distance of 1.2 nm, solvated with TIP3P water, neutralized and adjusted to 0.15 M NaCl. Proteins were parameterized using the CHARMM36 force field, and acyl-CoA parameters were generated using CGenFF. Residue protonation states were assigned at pH 7.4 using the H++ server, with the catalytic histidine modelled in the Nε2 tautomer. Simulations were performed using GROMACS 2024.2 on NVIDIA V100 GPUs. After 50,000 steps of steepest-descent minimization, each system was equilibrated for 100 ps under NVT conditions at 300 K using the V-rescale thermostat, followed by 100 ps under NPT conditions at 1 bar using the Parrinello–Rahman barostat. Production simulations were run for 200 ns with a 2 fs time step, and two independent replicas were performed for each system. Particle-mesh Ewald electrostatics, 1.2 nm electrostatic and van der Waals cutoffs, and LINCS bond constraints were applied.

Periodic-boundary artefacts were removed using sequential whole-molecule reconstruction, no-jump correction and centring. Catalytic competence was assessed from the Cys(Sγ)–acyl-C distance, using 4.0 Å as the productive threshold, the Bürgi–Dunitz nucleophilic-attack angle relative to the ideal value of approximately 107°^55^, and the geometry of the catalytic Cys–His–Asn triad. Substrate-pocket dimensions were monitored using distances between the Phe-gate and Phe-bottom residues, defined as GS Phe211/Phe270 and CURS Phe206/Phe265. Structural stability was evaluated from backbone RMSD, per-residue RMSF over the ordered protein core and ligand RMSD. Time series were smoothed using a 101-frame sliding window, and one- and two-dimensional density distributions were used to distinguish productive binding states from non-productive distance-trap conformations.

### Genome evolution and synteny analysis

Genome evolution analyses were performed using the haplotype-separated banana and ginger genome assemblies. For within-genome analysis, all-versus-all protein homology searches were conducted using DIAMOND. Candidate homologous gene pairs were integrated with chromosome position, strand orientation and gene order extracted from the corresponding GFF3 annotations. Collinear blocks were identified using the JCVI comparative genomics toolkit, and gene duplication modes were assigned according to collinear relationships, physical distance and local gene arrangement.

For comparative analyses among banana, ginger and turmeric, pairwise protein homology searches were performed using DIAMOND. The three highest-scoring homologues for each query were retained, including reciprocal best hits and additional high-confidence non-reciprocal matches, with reciprocal-best-hit status recorded for each pair. Homology results were combined with genomic coordinates and gene order information and used for interspecific collinearity analysis and anchor-pair identification using JCVI. Collinear blocks containing the target genes were subsequently extracted to evaluate their conservation, genomic context and evolutionary relationships across species.

### Quantification and statistical analysis

Data were processed in Excel or GraphPad Prism. The graphs represented means ± SD (n=3) unless otherwise indicated, as described in the figure legends. No statistical method was used to predetermine sample size. No data were excluded from the analysis. The experiments were not randomized. The investigators were not blinded to allocation during experiments and outcome assessment.

## Data availability

Source data are provided with this paper. The bulk RNA-seq data generated in this study have been deposited in the NCBI database under BioProject accession code PRJNA1503072 [https://www.ncbi.nlm.nih.gov/bioproject/PRJNA1503072]. The single-cell RNA-seq data generated in this study have been deposited in the NCBI database under BioProject accession code PRJNA1503074 [https://www.ncbi.nlm.nih.gov/bioproject/PRJNA1503074]. All other data supporting the findings of this study are available from the corresponding author upon reasonable request. The request will be responded within 2 weeks.

## Acknowledgements

This work was supported by the National Key Research and Development Program of China (2025YFA0921800 to B.L. and B.H.) and the Natural Science Foundation of China (22138006 to C.L., 22278240 to L.Q., 22478031 to B.L.).

## Author contributions

C.L., L.Q., B.L. and B.H. conceived the project. W.W. contributed to experimental design, planning, and execution. Q.L. analyzed the transcriptome data, J.Z. carried out enzyme structural modeling and molecular dynamics, X.Y. and X.L. helped in enzyme screening, F.M. developed the theoretical model. All authors contributed to data analysis and interpretation of results. W.W. drafted the manuscript. L.Q., B.L., B.H. and C.L. revised the manuscript and supervised the study.

